# Geometric morphometrics enables accurate predictions of paleoecology and reveals unique adaptations to an expanded niche space in extinct waterfowl

**DOI:** 10.1101/2025.05.29.656821

**Authors:** Ray Mariano Chatterji, Shante Williams, Janet C. Buckner

## Abstract

Establishing the relationships between organismal phenotypes and their environment is a key component to understanding evolutionary history. Comparative evaluations of extant and extinct species can reveal how lineages have adapted to changing environmental conditions over time. However, the understanding of paleoecologies is predicated on a robust understanding of how modern species have been shaped by adaptation. Waterfowl may present an ideal group to study adaptive evolution as much of their morphology is apparently shaped by their dietary ecology. Here we use a large geometric morphometric dataset of waterfowl combined with random forest, a supervised machine learning algorithm, and linear discriminate analysis, to predict the dietary ecologies of nine extinct waterfowl species. We find that both model types reliably predict ecology for extinct species with well-established diets. Interestingly, we also found that the Hawaiian moa-nalo and the New Zealand *Cnemiornis calcitrans* likely occupied ecological niches no longer present in modern waterfowl as they were not morphologically or ecologically convergent with modern geese as previously asserted. Our study demonstrates that waterfowl are an excellent model group for the study of adaptive evolution, and underscores the utility of predictive modelling for paleontological studies.

## INTRODUCTION

Understanding adaptive processes is a critical focus in the field of evolutionary biology and is fundamental to understanding several aspects of organismal biology and diversification (Reeve and Sherman 1993; Hoekstra and Coyne 2007; Wainright 2007; Losos 2011). Although research on adaptation over macroevolutionary timescales can be enriched through incorporation of paleoecological knowledge, direct evidence for the ecology of fossil species is often lacking. We can address this gap by leveraging investigations of adaptive morphological evolution in modern species, which can provide a baseline of comparison to interpret how selection has shaped the phenotypic diversity of extinct species, and further contributed to driving macroevolutionary patterns over the history of life (Kidwell and Flessa 2002; Hunt and Rabosky 2014; Fleagle et al. 2016).

Waterfowl (Anseriformes) present an excellent group to study morphological adaptation. Multiple studies have demonstrated that the skeletal anatomy of waterfowl is strongly influenced by foraging behaviour and diet (McCracken et al. 1999, Li and Clarke 2016, Olsen 2017, Chatterji et al. 2024). Repeated independent shifts into different dietary ecologies by various clades has resulted in convergent morphologies across multiple skeletal elements (McKracken et al.1999; Olsen 2017; Chatterji et al. 2024). This pattern of convergence in dietary ecology and associated morphologies extends to the waterfowl fossil record (Li and Clarke 2016; Olsen 2017; Buckner et al. 2018; De Mendozza and Gomez 2023). Previous attempts to infer the ecology of extinct waterfowl were limited to comparative morphology of a small number of species or morphometrics of single skeletal elements (Li and Clarke 2016; Olsen 2017; de Mendoza and Gomez 2021). However, Chatterji et al. (2024) showed that characters absent from previous studies, such as craniofacial angle, were important in distinguishing between different dietary ecologies. Further, the significant morphological integration between cranial and hindlimb shape recovered in Chatterji et al. (2024) suggests that comparative analyses using multiple elements should more accurately predict dietary ecology.

In this study, we use the Chatterji et al. (2024) dataset to predict the ecology of nine species of extinct waterfowl. Given the strong adaptive signal between diet and morphology found in that study, we hypothesized that skeletal shape in extinct species would reflect dietary adaptations. We first quantified the shape of the skull and hindlimb elements of nine recently extinct waterfowl species using geometric morphometrics. We then used random forest and linear discriminate analysis, trained on the dataset of modern species, to predict the diet and foraging behaviour of the extinct taxa. Our models reliably predicted the paleoecology of species which occupied similar morphospace to extant species. However, we found that the flightless species of Hawaii and New Zealand may display morphological adaptations to dietary niches which no longer exist among modern waterfowl.

## METHODS

Data Collection – Our dataset is modified from the morphometric and ecological dataset from Chatterji et al. (2024). We use the same set of landmarks for the skull (46 fixed landmarks, 313 semi landmarks in 21 curves), femur (17 fixed landmarks, 242 semi landmarks in 9 curves), tibiotarsus (8 fixed landmarks, 125 semi landmarks in 5 curves), and tarsometatarsus (13 fixed landmarks, 222 semi landmarks in 9 curves) to characterize shape in nine species of extinct anseriforms for all available elements (Table 1, S Table 1). Landmarks were placed in Slicer (Kikinis 2014) using the left limb elements, but if only right limb elements were available, they were mirrored in Meshlab (Cigoni et al. 2008). The cranial material of *Branta rhuax* was disarticulated, so we reconstructed it in Mehslab with refence to previous reconstructions (Paxinos et al. 2002) and other *Branta* specimens.

**Table 1:** Fossil specimens, their elements used in this study, and their ecotype as reported in the literature.

| Species | Elements | Reported ecotype | Dietary evidence |
| --- | --- | --- | --- |
| *Branta hylobadistes* | Femur, Tibiotarsus, Tarsometatarsus | Terrestrial Herbivore<sup>1,2</sup> | Morphology, Stable isotopes, closest living relatives |
| *Branta rhuax* | Skull, Femur, Tibiotarsus, Tarsometatarsus | Terrestrial Herbivore<sup>1,2</sup> | Morphology, Stable isotopes, closest living relatives |
| *Chendytes lawi* | Skull, Femur, Tibiotarsus, Tarsometatarsus | Diving Macroinvertevore<sup>3,4</sup> | Morphology, Stable isotopes |
| *Cnemiornis calcitrans* | Skull, Femur, Tibiotarsus, Tarsometatarsus | Terrestrial Herbivore<sup>5,6</sup> | Morphology, Stable isotopes, closest living relatives |
| *Mareca marecula* | Femur, Tibiotarsus, Tarsometatarsus | Unknown |  |
| *Ptaiochen pau* | Skull, Femur, Tibiotarsus, Tarsometatarsus | Terrestrial Herbivore<sup>1</sup> | Morphology |
| *Talpanas lippa* | Tibiotarsus, Tarsometatarsus | Unknown |  |
| *Thambetochen chauliodous* | Skull, Femur, Tibiotarsus, Tarsometatarsus | Terrestrial Herbivore<sup>1,7</sup> | Morphology, coprolites |
| *Thambetochen xanion* | Femur, Tibiotarsus, Tarsometatarsus | Terrestrial Herbivore<sup>1</sup> | Morphology |

Most of these species have uncertain phylogenetic placements. *Chendytes lawi* has been the most confidently placed using mitogenomic data (Buckner et al. 2018). *Thambetochen* and *Ptaiochen pau* have been tentatively placed as sister to the anatine (Sorenson et al. 1998) using fragmentary mitochondrial data. *Cnemiornis* has been placed as sister to the Australian *Cereopsis* based on morphological and molecular data (Worthy et al. 1997; Greer 2024). *Mareca marecula* and *T. lippa* have not been included in any phylogenetic analysis and their phylogenetic affinity is unknown. Due to the uncertainty, which can be common for fossil species, we did not use tree-based methods such as phylogenetic flexible discriminate analysis (Motani and Schmitz 2011).

We followed the protocol from Chatterji et al. (2024) to ensure continuity of the dataset. All statistical analysis was performed in R 4.4.3 (R Core Team 2021). Missing data was estimated using the geomorph v4.0 (Collyer and Adams 2018) function *estimate*.*missing*. Curves were resampled and semilandmarks placed equidistant along them using the SURGE (Felice 2019) function *subsampl*.*inter*. All data was then subject to Procrustes superimposition using the geomorph function *gpagen*. We ran a general linear model comparing shape and size for each element using the geomorph function *procD*.*lm* to test for the effects of allometry. We then removed and separately analyzed the size free residuals from the general linear model to test whether the models performed better with size removed. We then quantified and visualised shape variation among elements using a principal component analysis (PCA) with the geomorph function *gm*.*prcomp*. Significant PCs (≥5% of variation explained) were used in the subsequent discriminate analyses.

We chose fossil species based on availability of specimens, and most of the taxa have dietary ecologies reported with reasonable supporting evidence (Table 1). These independent sources of evidence make them ideal species to test how well morphometric data can predict the dietary ecology of extinct taxa. We also include *Mareca marecula* (Olson and Jouventin 1995) and *Talpanas lippa* (Iwanuik et al. 2009) which have uncertain dietary ecologies to make naïve predictions about poorly known taxa. Ecology was broken into three categories: diet, the most common food type for a species; foraging, the dominant method for obtaining food; and ecotype, a combination of diet and foraging. Assignments of ecological categories for modern taxa is taken form Chatterji et al. (2024).

Predictive models – We used two methods modified from Sosiak and Barden (2020) for ecological niche prediction, multi group linear discriminate analysis (LDA) and random forest (RF). The LDAs were performed using the MASS (Venables and Ripley 2002) function *lda*. This method identifies between group variation and maximizes this variation by reducing the dimensionality of the data to the most informative variables. The fossil species are then assigned to their most likely group based on trait values distance to the group mean of the specified ecological category. RF methods (R package RandomForest; Liaw and Wiener, 2018) are a supervised machine learning program based on a series of decision trees which are used to construct a ‘consensus tree’. Each model used ntree = 5000. We set the mtry for each element, the number of variables to be removed each iteration, to 1/3 of total model variables or the closest fraction.

We created an LDA and a RF model for each of the four skeletal elements independently to predict diet, foraging, and ecotype state for the extinct species. We then combined all significant PCs across all elements to create an LDA and a RF model for species with complete data (all four elements; Table 1), for a total of 30 models. We tested the accuracy of each model by training it on a random 80% subset of the extant dataset, then tested its predictive accuracy on the remaining 20%. We performed 20 such tests for each model and rejected models with greater than 30% error rate. We then repeated all analyses for the size free datasets.

## RESULTS

### Fossil taxa extend the known morphospace for waterfowl

All elements were significantly related to size (P<0.005). How much size contributed to shape varied between elements, from 16.8% in the femur to 4.5% for the tibiotarsus. PCAs did not change qualitatively with size free data (S Fig 1).

While the addition of fossils did not change the major axes of variation described by PC1 and PC2 for each of the elements (Chatterji et al., 2024), they extended them. The moa-nalo (*Thambetochen spp*. and *Ptaiochen pau*) and *Cnemiornis calcitrans* exhibit extreme values of PC1 or PC2 for each element (Fig 1). For the skull (Fig 1A), PC1 broadly describes proportional bill length and skull height, while PC2 largely describes craniofacial angle. PC1 for the femur (Fig 1B) represents the angle of the trochlea and the size of the femoral trochanter, while PC2 describes the size of muscular scars and the twisting of the shaft. For the tibiotarsus (Fig 1C), PC1 largely describes the proximal projection of the cnemial crest, and PC2 describes the width of the proximal end as well as the anterior projection of the cnemial crest. PC1 for the tarsometatarsus (Fig 1D) describes its relative elongation, and PC2 largely describes the width of the troachlea, the placement of trochlea II, and the length of the crista hypotarsi.

**Figure 1.**
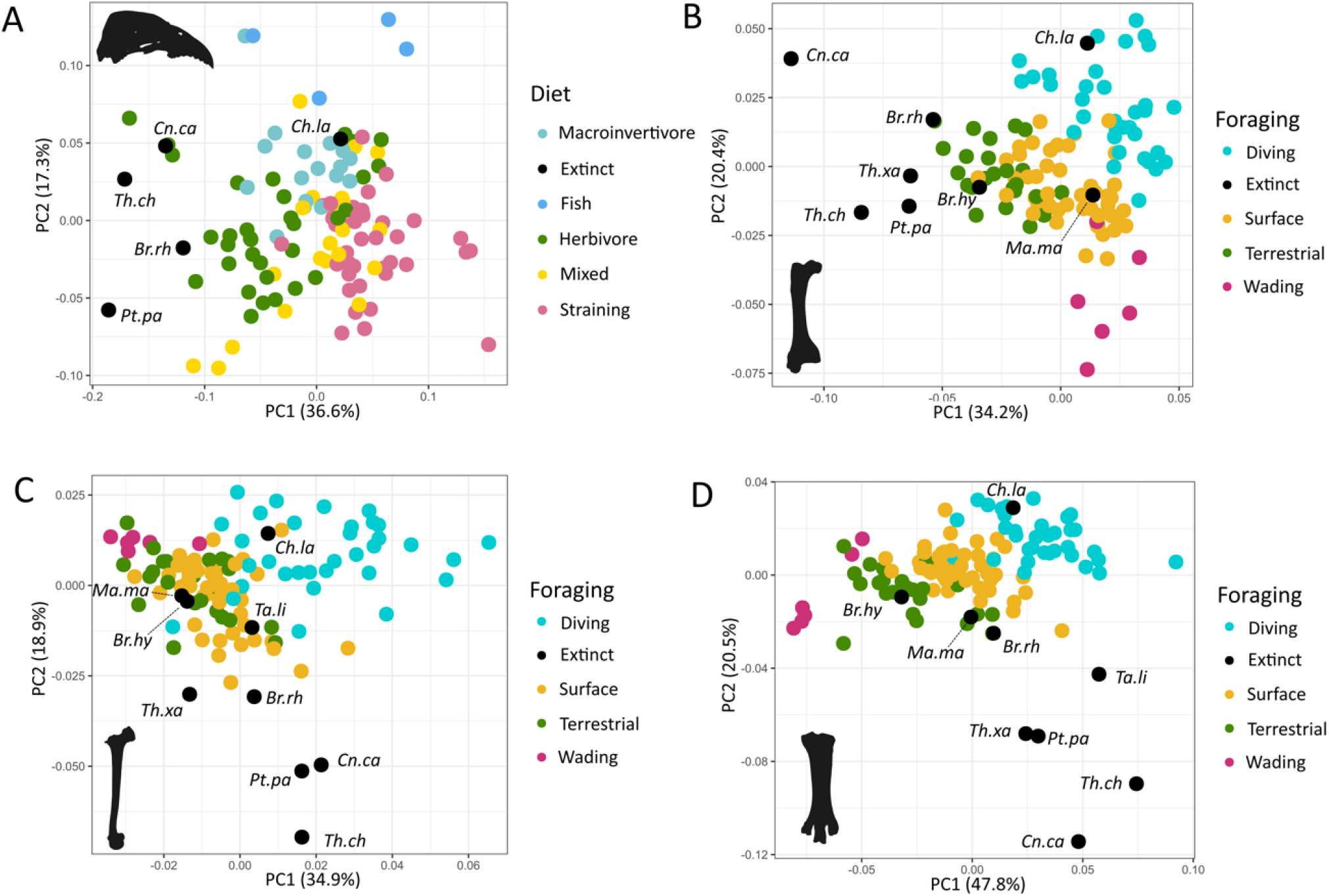
PCAs for each of the four elements for PC1 and PC2: skull (A), femur (B), tibiotarsus (C), tarsometatarsus. Species coloured by ecological category with the highest predictive accuracy based on the RF and LDA models.

*Thambetochen spp*., *Ptaiochen pau*, and *Cnemiornis calcitrans* consistently exhibiti the most negative values in key parts of the morphospace of each element. In the skull these species have shorter bills and taller, more rounded skulls than other taxa (Fig. 1A). The femur of all four species has a larger femoral trochanter, and wider distal end that is twisted medially (Fig. 1B), with particularly extreme values for *Cn. calcitrans. Th. chauliodous, P. pau*. and *Cn. calcitrans* also had much wider tibiotarsi, particularly the proximal portion, and a larger anteriorly projecting cnemial crest (Fig. 1C). These species’ tarsometatarsi (Fig 1D) are characterized by a broader distal region, with more splayed trochlea. Specifically, trochlea II is more distally located and the medial crista hypotarsi is longer. *Talpanas lippa* also has a lower tarsometatarsus PC 2 value than other waterfowl, excluding the previously mentioned fossil species.

### A Random Forest model trained on skull and hindlimb shape best predicts dietary ecology of extant species

Our predictive models varied in accuracy based on the element, the ecological variable, and the method. In all cases the size corrected data models were less accurate than their counterparts (S Table 2). We will therefore focus on the results for models including size. After rejecting models with > 30% error rate, we had a final total of 13 models consisting of parallel sets of RF and LDA models (Table 2).

**Table 2:** Single element and combined models meeting error rate cut off.

| **Element** | **Model type** | **Category** | **Error rate mean** |
| --- | --- | --- | --- |
| Combined | Random Forest | Foraging | 9.8% |
| Combined | LDA | Foraging | 11.2% |
| Tarsometatarsus | Random Forest | Foraging | 17.7% |
| Femur | Random Forest | Foraging | 18.9% |
| Combined | Random Forest | Ecotype | 19.3% |
| Tarsometatarsus | LDA | Foraging | 20.3% |
| Combined | LDA | Diet | 20.3% |
| Combined | Random Forest | Diet | 21.3% |
| Femur | LDA | Foraging | 22% |
| Skull | Random Forest | Diet | 22.5% |
| Skull | LDA | Diet | 27.2% |
| Skull | Random Forest | Foraging | 28.3% |
| Combined | LDA | Ecotype | 29.8% |

The combined-RF model, based on all shape data from the skull and hindlimb, produced the highest predictive accuracy for foraging and ecotype (Table 2). Element-specific RF models each successfully predicted only one ecological category, except for the skull which predicted both diet and foraging. The combined-LDA models have near equivalent predictive accuracy to their RF counterparts (Table 2), except the combined-diet model, with a lower predictive accuracy of 70%.

### Model predictions support reported dietary ecologies for fossil taxa that fall within extant morphospace, but perform poorly on extinct outliers

Ecological predictions were generally consistent with previous hypotheses for species that fell within the extant morphospace. The tibiotarsus had the lowest predictive accuracy given a high degree of morphospace overlap among non-diving species, so we excluded it from our ecological assessment of fossil taxa. Extinct *Branta* species were predicted as terrestrial based on hindlimb shape, apart from the femur-foraging RF model that predicted *B. hylobadistes* was a surface swimmer (Fig. 2C). Skull-diet models predicted an herbivorous diet for *B. rhuax* (Fig 2A-B S Table 2). All hindlimb-foraging models predicted that *Ch. lawi* was a diving species (Fig. 2C-F, S Table 4-5). The skull-diet models predicted an herbivorous diet (Fig. 2A-B, S Table 3), contradicting previous morphological and isotopic analyses, likely due to missing data resulting from the incomplete skull. However, combined RF and LDA (Fig. 3, S Table 6-7) models predicted a macroinvertivore diet along with a macroinvertivore/diving ecotype.

**Figure 2.**
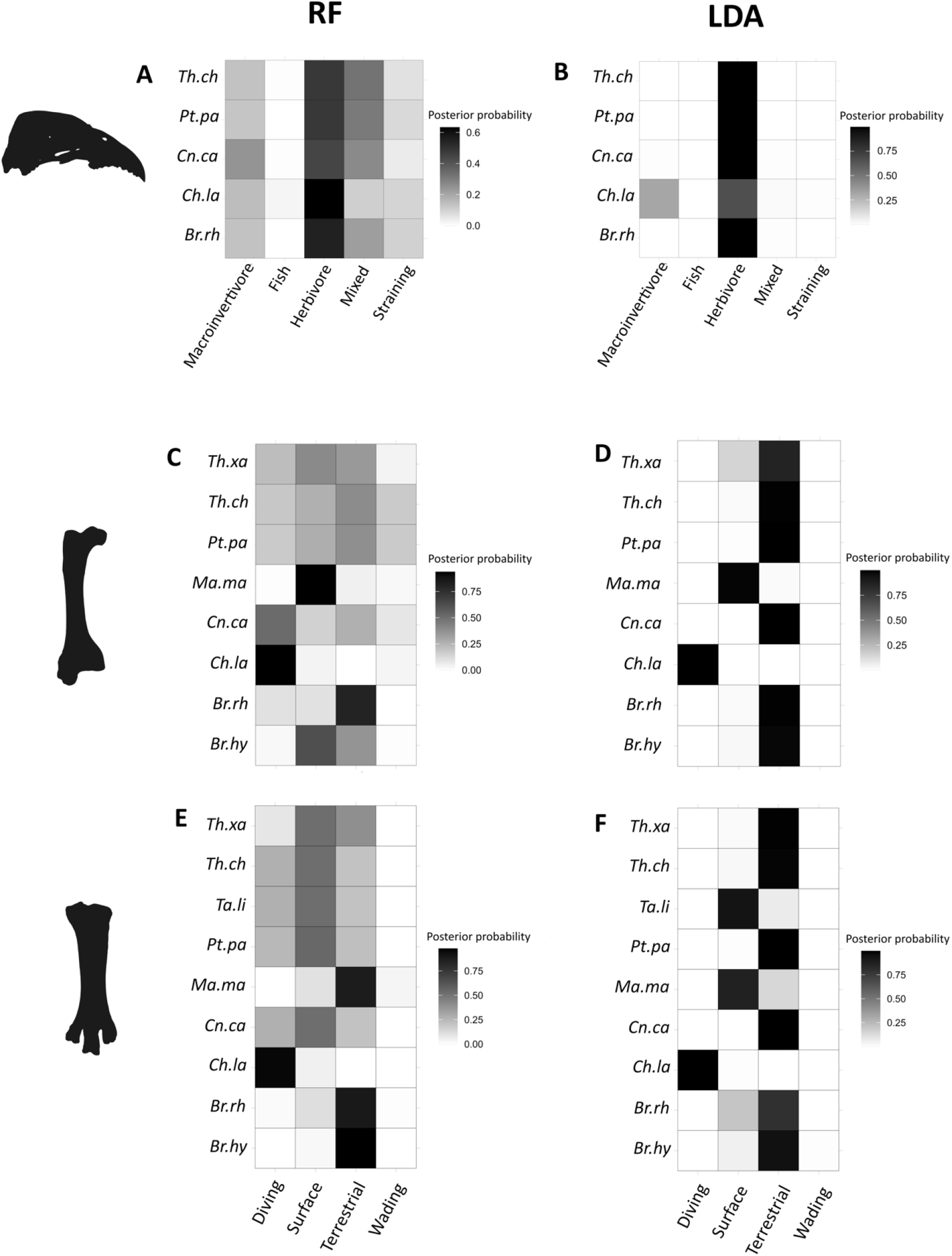
Posterior probability values of the most accurate models for each element for RF (left) and LDA (Right) excluding the tibiotarsus which had no models with error rates lower than 30%. A-B skull, C-D femur, E-F tarsometatarsus.

**Figure 3.**
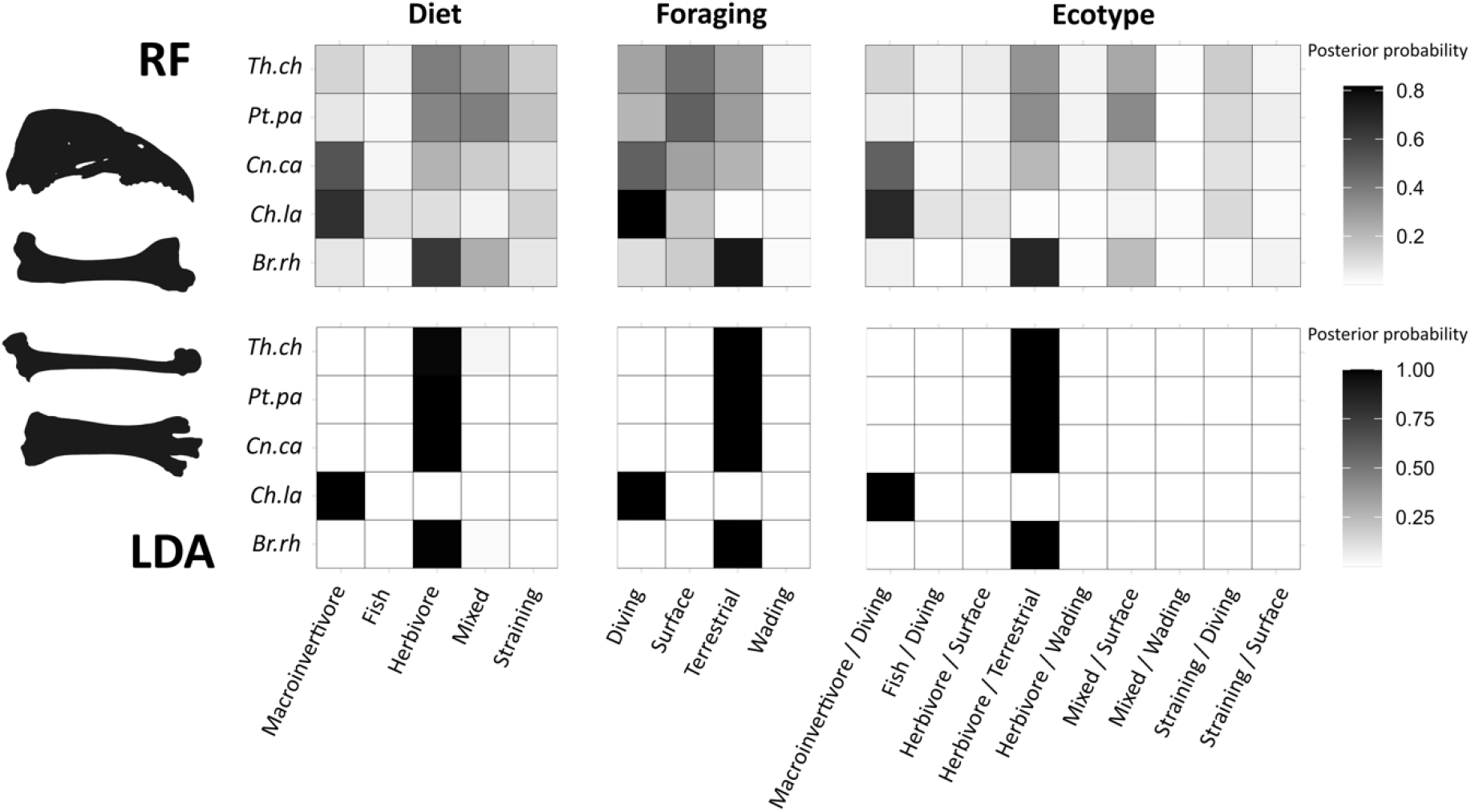
Posterior probability values of RF (top) and LDA (bottom) combined model predictions.

The models predicted variable ecological states for the morphological outliers, moa-nalo and *Cn. calcitrans*. The skull-diet RF and LDA (Fig. 2A-B, S Table 3) models predicted an herbivorous diet, as expected based on previous research. The LDA femur-foraging model (Fig 2.D, S Table 4) predicted that they were all terrestrial. However, the foraging RF (Fig. 2C, S Table 4) models produced a range of results that were poorly supported. There was disagreement between tarsometatarsus-foraging models (Fig. 2E-F, S Table 5) with the RF model predicting surface swimming for all four species while the LDA model predicted terrestrial for all four species. The combined-ecotype RF model (Fig 3, S Table 6) made weakly supported predictions for these species: *Th. chauliodous* – herbivorous/terrestrial ecotype (pp = 0.31), *P. pau* – mixed/surface ecotype (pp = 0.42), and *Cn. calcitrans* – macroinvertivore/diver (pp= 0.46). The combined-ecotype LDA model (Fig 3; S Table 7) predicted all three species to be herbivore/terrestrial (pp > 0.7), which best aligns with previous hypotheses.

For two species, we have no reliable dietary data, meaning our predictions are naïve. *Mareca marecula* was predicted to be a surface swimmer by the femur-foraging RF and LDA models (S Table X), consistent with its congeners. The tarsometatarsus-foraging LDA Model predicted it to be a surface swimmer, while the corresponding RF model predicted it to be terrestrial (S Table 5). The tarsometatarsus-foraging models predicted *Talpanas lippa* to be a surface swimmer (S Table 5), but this taxon also falls outside of extant morphospace (Fig. 1).

## DISCUSSION

### Predictive models are useful tools for morphometrics-based investigations of paleoecology

Here we use geometric morphometric data to develop predictive models for classifying the ecologies of extinct waterfowl species. The success of these models supports the long history of using morphology to describe ecology of fossil waterfowl. Our combined models, that jointly considered all available morphometric data, were the most accurate and assigned extinct species to their reported diet and foraging categories. The high accuracy of the combined-foraging models corroborates Chatterji et al. (2024) finding that foraging exerts the greatest selective pressure on hind limb and skull shape in waterfowl. Further, the universally more accurate predictions of the combined models support that the hindlimb and skull are evolutionarily integrated with one another. In this sense, making predictions based on multiple morphological characters clearly increases predictive power. However, multiple extinct species occupied areas of the morphospace not represented in modern species, limiting model performance due to lack of appropriate training data (see below).

Previous palaeoecological assessments of waterfowl morphology were based on qualitative descriptions (Olson and James 1991; Worthy et al. 1997; Zelenkov 2020) or were not model based (Li and Clarke 2016; Olsen 2017). Mendoza and Gomez (2021) is the only study applying a phylogenetically flexible discriminate analysis to predict foraging ecology based on the tarsometatarsus. Our tarsometatarsus-foraging models performed similarly and accurately predict foraging ecology. However, our multi-element approach allowed us to accurately predict more complex ecological categories such as ecotype. In this sense, we confirm that evaluating multiple morphological elements clearly increases predictive power and sensitivity.

These ecomorphological predictive models can be powerful tools for paleontological research (Sosiak et al. 2023) but have limitations. The model types were comparable in accuracy, with RF models being slightly more accurate and LDA models generally having higher support values. LDA models also predicted ecologies more in line with our previous knowledge of the extinct species. However, LDA reduces data dimensionality which likely explains its very high support values for all predictions, despite the distinct shapes of many skeletal elements across fossil taxa. The low support values of RF models may reflect unique shapes of the fossil taxa but consequently poorly predicted their ecology. Both models also rely on predesignated categories which is problematic for evaluating extinct species with unobserved ecologies, as discussed in detail below (Godoy 2020; Varela et al. 2023).

### Evaluations of fossil morphometrics extend the known morphospace for waterfowl and provides insights into their paleoecology and phenotypic evolution

The models reliably predicted the paleoecology of fossil species that fall within extant morphospace. *Chendytes lawi* was originally described as a large, flightless sea duck (tribe Mergini) from coastal and insular California. Molecular systematics later showed that the morphological similarities between *C. lawi* and sea duck species resulted from convergence, most likely related to diet and foraging. Recovery of *C. lawi* as a macroinvertivore diver is consistent with previous paleoecological hypotheses that suggested it specialized on sessile marine invertebrates (Miller et al., 1961; Livezey, 1993). Later, dietary isotopic analysis reinforced this hypothesis by revealing isotopic signatures consistent with the largely carnivorous diets of harbor seals and sea otters (Jones et al., 2021).

The models also predicted that the two extinct Hawaiian *Branta* species were terrestrial herbivores, consistent with their extant congeners (James and Olson 1991; Paxinos et al. 2001). All modern terrestrial herbivorous waterfowl have a remarkably consistent morphology (Chatterji et al. 2024), as seen in Figure 1. However, some fossil species are notably divergent, occupying distinct areas of this morphospace (Fig. 1,4). The moa-nalo, generally described as convergent with geese (James and Olson 1991; Olsen 2017), have comparatively short, curved bills, broader femurs, more robust tibiotarsi, and stouter tarsometatarsi with widely splayed trochlea. Previous research suggested that their short, curved bill and pseudodentition are indicative of a specialist browsing diet (James and Burnery 2008). The limited distributional overlap between *Th. chauliodous* and *P. pau* may be explained by their occupation of a shared or similar browsing niche as is suggested by their similar morphologies (James and Olson 1991). Coprolite data from *Th. chauliodous* revealed significant consumption of ferns as well as evidence for hindgut fermentation, both suggesting a distinct feeding strategy from modern geese (James and Burney 2008). Thus, browsing specialization may represent a moa-nalo synapomorphy.

**Figure 4.**
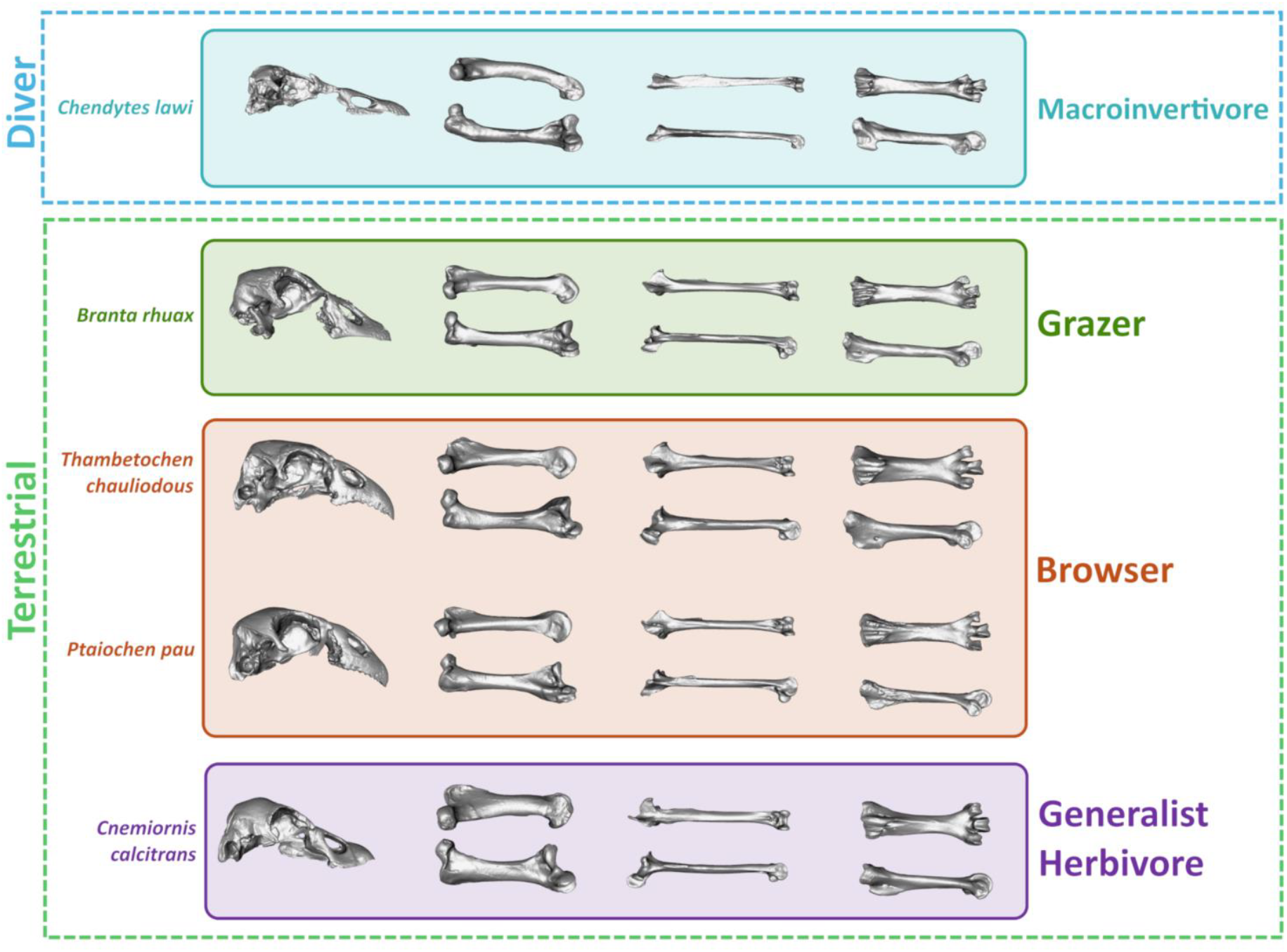
The proposed ecological niches of the five fossil species for which we analysed all four elements.

The morphological divergences between browsers and grazers are well-studied in mammalian herbivores (Clauss et al. 2008; Mitchell et al. 2018; Codoron et al. 2019) but remains underexplored in birds. In mammals, browsers and herbivores exhibit distinct morphologies, such as tooth shape, limb length, and craniofacial shape (Clauss et al. 2008). Though comparatively understudied, notable cranial similarities among browsing birds (moa, hoatzin, moa-nalo) include a robust downward curving bill which provides a stronger bite force and is likely an adaptation to cropping tough fibrous or woody material (Attard et al. 2016). Contrarily, most grazing waterfowl have short but comparatively gracile, straight beaks (Olsen 2017; Chatterji et al. 2024). This dichotomy is similar to macropodid morphology (kangaroos and relatives), where craniofacial shape is associated with the biomechanics of acquiring the food: shorter stronger skulls are associated with cropping and longer more gracile skulls are associated with plucking (Mitchell et al. 2018). Most grazing waterfowl forage by biting and then plucking grass rather than cropping (Kurk 2008), while the bill curvature and pseudodentition of the moa-nalo suggest that they cropped their food.

*Cnemiornis calcitrans* has previously been classified as a specialist grazer due to its sister relationship with *Cereopsis novaehollandiae* and its short, wide, and squared bill (Worthy et al. 1997). However, we found that *Cn. calcitrans* consistently occupied distant areas of the morphospace from modern grazers. This is best illustrated by the contrast between *C. calcitrans’* short, stocky tarsometatarsi and the exceptionally elongate tarsometatarsi of its sister genus *Ce. novaehollandiae*. The limbs of *Cn. calcitrans* are more like the browsing, forest dwelling moa-nalo. Stable isotope analysis revealed ^15^N levels that indicated herbivory, but distinct ^13^C values that potentially indicate niche partitioning among herbivorous anatid species (Wood et al., 2017). Radiocarbon dated specimens, along with extensive pollen records, suggest that *Cn. calcitrans* was more varied in its habitat than originally believed (Johnston et al. 2022). Evidence that they occurred in dense woodlands may mean that its stouter and more robust limbs were an adaptation to generalized use of a variety of habitats including grasslands, sedge swamps and woodland.

Our data underscores the unique features of hind limb morphology in forest dwelling species. Though *Cn. calcitrans* may be found in both open and closed environments, moa-nalo specimens are found largely in forested environments (Olson and James 1991; James and Burney 2008). All hind limb bones, particularly the tibiotarsus and tarsometatarsus, were comparatively more robust in the forest dwelling species than in grazers (Fig 3). A parallel pattern exists between the plains-dwelling emu *(Dromaius novaehollandiae)* and the forest-dwelling cassowary (*Casuarius casuarius*) (Fowler 1991); and between hooved mammals living in open versus closed environments (Clauss et al. 2008). Presumably, open environments select for longer more efficient strides, while more closed environments select for shorter, more stable limbs (Curran 2012; Etienne et al. 2020).

The moa species of New Zealand provide an interesting evolutionary parallel. This group of terrestrial birds was spread across multiple biomes and display a wide range of hindlimb morphology. Moas achieved varying levels of cursoriality (e.g., long legged *Dinornis*) and robustness (e.g., *Pachyornis*) (Kooyman 1991; Worthy and Scofield 2012; Brassey et al. 2013). *Pachyornis* had a significantly more stress resistant tibiotarsus and femur than more cursorial ratites (Brassey et al. 2013). Though not tested specifically it is notable that the more robust species of moa have noticeably shorter and stockier tarsometatarsi, similar in proportion to the moa-nalo and *Cn. calcitrans*, presumably advantageous for stress resistance and balance. The ecological significance of this hindlimb variation among moa is currently unknown as both morphs are largely sympatric with broad dietary overlap (Worthy and Schofield 2011). A comprehensive analysis of hindlimb evolution in terrestrial birds could shed light on the sources of this variation.

### Potential insights on paleoecology from naïve model predictions

Our models may provide valuable insight on poorly studied extinct taxa that fall within extant morphospace. *M. marecula* was a flightless species endemic to Amsterdam Island, presumably driven to extinction by hunting and introduced rats (Bourne et al. 1983). Most of our models predicted *M. marecula* be a surface swimmer, consistent with its *Mareca* and *Anas* relatives, most of which are surface swimming species. However, the island’s limited fresh water, *M. mareca’s* lack of developed salt glands, its relatively robust hindlimb, and frequent recovery of fossils from the interior of the island were together interpreted as evidence for significant terrestrial habit rather than coastal (Olson and Jouventin 1995). The tarsometatarsus-foraging RF model and the PCA (Fig 2E) of the tarsometatarsus does suggest some adaptation towards increased terrestriality (pp=0.87). This prediction might hold additional weight as we found that the tarsometatarsus had the highest predictive power for foraging type. Recent divergence from a morphologically typical population of dabbling ducks may explain a lack of more obvious terrestrial adaptations in other elements. However, among waterfowl, a pattern of island colonization followed by a rapid loss of flight, and increased terrestrially is not uncommon (Worthy 1998; Sorenson et al. 1999; Rosinger et al. 2024). A more detailed morphometric study of the few dozen individual fossil specimens, including the skull which we were unable to measure, may provide more clarity on this interpretation.

*Talpanas lippa* is another extinct, flightless island endemic from Hawaii and may be one of the most morphologically distinct waterfowl species known. Given *T. lippa* was likely blind (small orbits and reduced visual sensory system) and exhibited a greatly expanded somatosensory system relative to other waterfowl, it may have been a nocturnal tactile-foraging invertivore specialist (Iwaniuk et al. 2009; Witmer et al. 2018). We were unable to measure the partial skull holotype, but tarsometatarsus models consistently predicted *T. lippa* to be a surface swimmer, with mixed support (RF pp = 0.52, LDA pp = 0.92). Given the lack of other elements, we are unable to confidently reconstruct its foraging ecology. We suspect that the unique features of the skull would likely place *T. lippa* as another fossil outlier in waterfowl morphospace, such that predictions based on modern diversity may not prove very useful. Like the other Hawaiian fossil species, it occupies a distinct part of tarsometatarsus morphospace, having a more robust tarsometatarsus, wider spread trochlea and distal trochlea II. Combined with its broad posterior pelvis and shallow cnemial crest, the morphology suggests a terrestrial foraging habit (Iwaniuk et al. 2009) and could indicate similarities to other forested species, selecting for stability rather than increased stride length (Clauss et al. 2008). If *T. lippa* was terrestrial, it adapted to this lifestyle through distinct means from modern terrestrial waterfowl. Such unique ecologies, frequently recovered for island endemics, which fall outside of the modern framework limits the effectiveness of predictive models. Future models may need to incorporate these distinctive ecologies into their parameters to increase accuracy.

## CONCLUSION

Here we find that geometric morphometric datasets can be used to correctly predict the ecology of extinct species if they fall within the sampled variation. Both random forest and linear discriminate analysis show great potential for ecomorphological analysis of paleontological datasets. However, we found that the models performed poorly if species fell far outside of the test data’s variation. The moa-nalo, *Cnemiornis calcitrans*, and *Talpnanas lippa* fell outside of the extant range of variation, suggesting that these species have adapted to niches unoccupied by extant waterfowl. Thus, including extinct species into a comprehensive quantitative dataset of modern taxa revealed novel insights into the adaptive morphology of waterfowl. Our work further reinforces the promise of waterfowl in adaptation studies, displaying an even wider diversity of ecologies and associated morphologies than previously thought.

## ACKNOWLEDGMENTS

We would like to thank Sean Mcleod and Helen James for access to their collections. We would also like to Helen James for thoughtful discussions and her knowledge about the ecologies and anatomy of the fossil species.

## Notes

### Competing Interest Statement

The authors have declared no competing interest.

