## Appendix 1 for "Geometric morphometrics enables accurate predictions of paleoecology and reveals unique adaptations to an expanded niche space in extinct waterfowl"

**Paleoecology references**

Supplementary table 2. Error rates of rejected models

| **TYPE** | **RF** | **LDA** |
| --- | --- | --- |
| Sk - Ecotype | 38.5 | 46.6 |
| Tarso _diet | 56.4 | 51 |
| Tarso_ecotype | 59.3 | 52.3 |
| Fem_diet | 39.6 | 35.7 |
| Fem_ecotype | 46.6 | 36.1 |
| Tib_for | 31.8 | 32.74 |
| Tib_Diet | 53.1 | 64.8 |
| Tib_Cat | 52.1 | 65.3 |
| SF-Sk - Ecotype | 43.8 | 38.7 |
| SF-Sk - Diet | 31.6 | 34.2 |
| SF-Sk - Foraging | 29.7 | 29.5 |
| SF-Tarso _diet | 67.3 | 53.7 |
| SF-Tarso_ecotype | 71.9 | 63.1 |
| SF-Tarso_Foraging | 31.7 | 32.7 |
| SF-Fem_diet | 38.7 | 52.5 |
| SF-Fem_ecotype | 41.6 | 47.5 |
| SF-Fem_Foraging | 21.6 | 24.1 |
| SF-Tib_for | 31.8 | 37.1 |
| SF-Tib_Diet | 52.2 | 61 |
| SF-Tib_Ecotype | 51.3 | 61.7 |

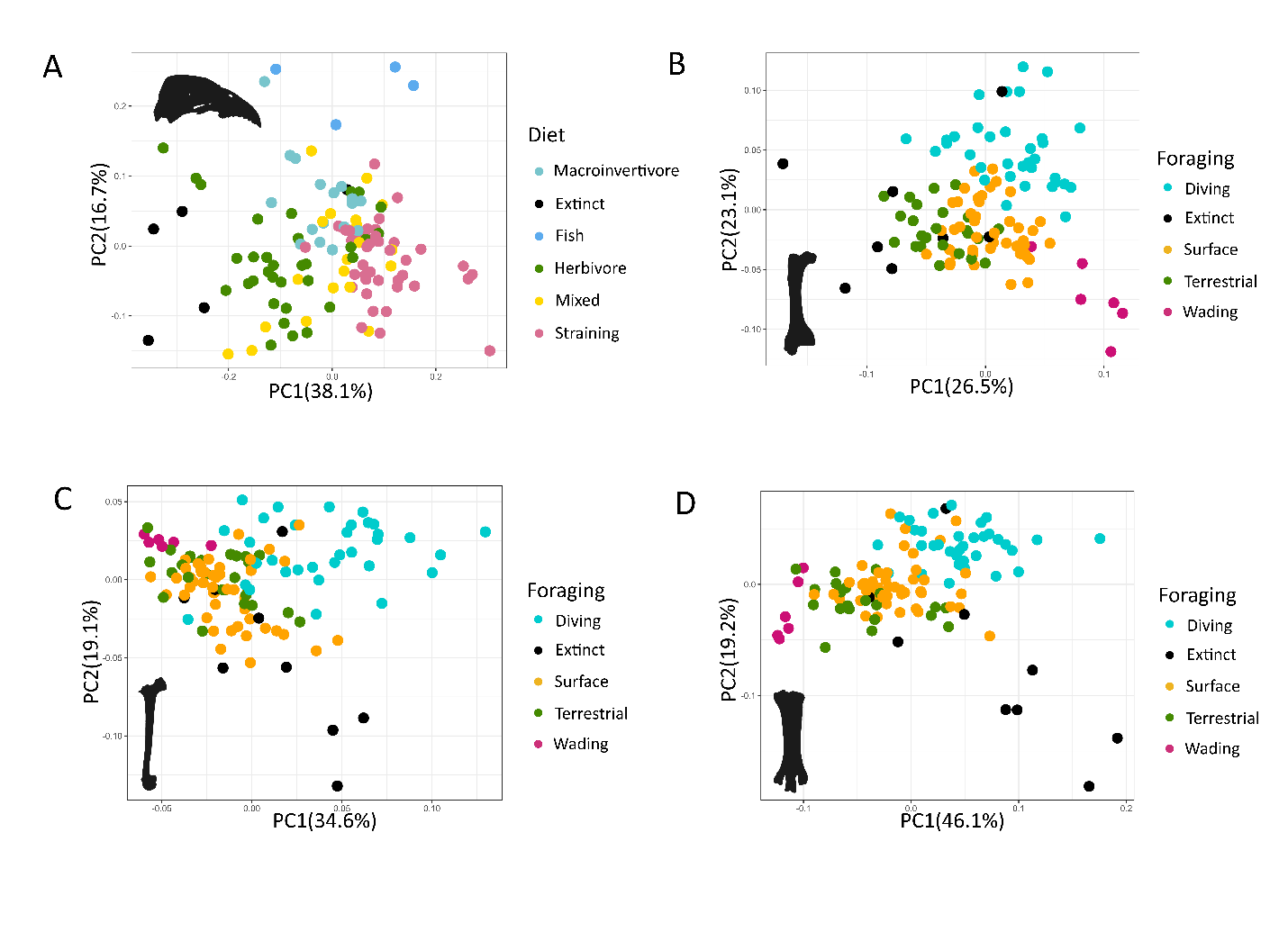

Supplementary Figure 1. Size Free PCAs for each of the four elements for PC1 and PC2: skull (A), femur (B), tibiotarsus (C), tarsometatarsus. Species coloured by ecological category with the highest predictive accuracy based on the RF and LDA models.

Supplementary table 3. Predictions of models and their posterior probabilities for the Skull-Diet models. LDA, linear discriminate analysis; RF, random forest; PP, posterior probability; SF, Size free.

| **Species** | **Random Forest** | **Random Forest PP** | **LDA** | **LDA PP** |
| --- | --- | --- | --- | --- |
| Branta rhuax | Herbivore | 0.54 | Herbivore | 0.99 |
| Chendytes lawi | Herbivore | 0.62 | Herbivore | 0.66 |
| Cnemiornis calcitrans | Herbivore | 0.47 | Herbivore | 0.99 |
| Ptaiochen pau | Herbivore | 0.48 | Herbivore | 0.99 |
| Thambetochen chauliodous | Herbivore | 0.49 | Herbivore | 0.99 |

Supplementary table 4. Predictions of models and their posterior probabilities for the Femur-Foraging models. LDA, linear discriminate analysis; RF, random forest; PP, posterior probability; SF, Size free.

| **Species** | **Random Forest** | **Random Forest PP** | **LDA** | **LDA PP** |
| --- | --- | --- | --- | --- |
| Branta hyblodites | Surface | 0.61 | Terrestrial | 0.97 |
| Branta rhuax | Terrestrial | 0.81 | Terrestrial | 0.99 |
| Chendytes lawi | Diving | 0.94 | Diving | 0.99 |
| Cnemiornis calcitrans | Diving | 0.51 | Terrestrial | 0.99 |
| Mareca marecula | Surface | 0.93 | Surface | 0.98 |
| Ptaiochen pau | Terrestrial | 0.37 | Terrestrial | 0.99 |
| Thambetochen chauliodous | Terrestrial | 0.38 | Terrestrial | 0.98 |
| Thambetochen xanion | Surface | 0.39 | Terrestrial | 0.85 |

Supplementary table 5. Predictions of models and their posterior probabilities for the Tarsometatarsus-Foraging models. LDA, linear discriminate analysis; RF, random forest; PP, posterior probability; SF, Size free.

| **Species** | **Random Forest** | **Random Forest PP** | **LDA** | **LDA PP** |
| --- | --- | --- | --- | --- |
| Branta hyblodites | Terrestrial | 0.72 | Surface | 0.66 |
| Branta rhuax | Surface | 0.53 | Surface | 0.92 |
| Chendytes lawi | Diving | 0.69 | Diving | 0.92 |
| Cnemiornis calcitrans | Diving | 0.62 | Diving | 0.97 |
| Mareca marecula | Surface | 0.93 | Surface | 0.64 |
| Ptaiochen pau | Diving | 0.49 | Surface | 0.96 |
| Talpanas lippa | Surface | 0.59 | Surface | 0.80 |
| Thambetochen chauliodous | Diving | 0.63 | Surface | 0.98 |
| Thambetochen xanion | Terrestrial | 0.38 | Surface | 0.86 |

Supplementary table 6. Predictions of models and their posterior probabilities for the combined random forest models. RF, random forest; PP, posterior probability; SF, size free.

| **Species** | **RF - Diet** | **RF - Diet**  **PP** | **RF - Foraging** | **RF - Foraging**  **PP** | **RF - Ecotype** | **RF - Ecotype**  **PP** |
| --- | --- | --- | --- | --- | --- | --- |
| Branta rhuax | Herbivore | 0.61 | Terrestrial | 0.77 | Herb/terr | 0.69 |
| Chendytes lawi | Marcoinvertivore | 0.67 | Diving | 0.81 | Macro/Dive | 0.67 |
| Cnemiornis calcitrans | Marcoinvertivore | 0.53 | Diving | 0.48 | Macro/Dive | 0.47 |
| Ptaiochen pau | Mixed | 0.38 | Surface | 0.47 | Mixed/surf | 0.35 |
| Thambetochen chauliodous | Herbivore | 0.37 | Surface | 0.42 | Herb/terr | 0.31 |

Supplementary table 7. Predictions of models and their posterior probabilities for the combined linear discriminate analysis models. LDA, linear discriminate analysis; PP, posterior probability; SF, size free.

| **Species** | **LDA - Diet** | **LDA - Diet**  **PP** | **LDA - Foraging** | **LDA - Foraging**  **PP** | **LDA - Ecotype** | **LDA - Ecotype**  **PP** |
| --- | --- | --- | --- | --- | --- | --- |
| Branta rhuax | Herbivore | 0.98 | Terrestrial | 0.99 | Herb/terr | 0.99 |
| Chendytes lawi | Marcoinvertivore | 0.99 | Diving | 0.99 | Macro/Dive | 0.99 |
| Cnemiornis calcitrans | Herbivore | 1 | Terrestrial | 1 | Herb/terr | 1 |
| Ptaiochen pau | Herbivore | 0.99 | Terrestrial | 1 | Herb/terr | 0.99 |
| Thambetochen chauliodous | Herbivore | 0.97 | Terrestrial | 0.99 | Herb/terr | 0.99 |
